# Purification of age-distinct insulin secretory granules through antigen restriction

**DOI:** 10.1101/2020.06.03.103770

**Authors:** Martin Neukam, Pia Sala, Andreas-David Brunner, Katharina Ganß, Alessandra Palladini, Michal Grzybek, Jovana Vasiljević, Johannes Broichhagen, Kai Johnsson, Thomas Kurth, Matthias Mann, Ünal Coskun, Michele Solimena

## Abstract

Endocrine cells employ regulated exocytosis of secretory granules to secrete hormones and neurotransmitters. The competence of secretory granules for exocytosis depends on spatio-temporal variables such as proximity to the plasma membrane and age, with newly-generated granules being preferentially released. Despite recent advances, we still lack a comprehensive view about the molecular composition of insulin granules and how this may change over the lifetime of the organelles. Here we report a strategy for the purification of insulin secretory granules of distinct age from insulinoma INS-1 cells. Tagging the granule-resident protein phogrin with a cleavable CLIP tag, we obtained intact fractions of age-distinct granules for proteomic and lipidomic analyses. We find that the lipid composition changes over time along with the physical properties of the membrane.

## Introduction

Exocytosis is a fundamental cellular process to transport cargoes out of cells and to maintain plasma membrane (PM) protein and lipid homeostasis. Coordinated secretion in time and space is further crucial for the maintenance of extracellular matrix and cell-cell communication. In addition to constitutive exocytosis, which is constantly occurring in all cells, specialized professionally secreting cells employ regulated exocytosis. Here, a specific trigger is required to elicit secretion of vesicles awaiting exocytosis upon elevation of intracellular Ca^2+^ levels. This pathway is utilized by many cell types, including neurons and peptide-hormone secreting endocrine cells. As a member of the latter group, pancreatic β-cells require stimulation by glucose or other secretagogues to secrete insulin.

Similar to other cargoes undergoing regulated exocytosis, insulin is produced as an inactive prohormone precursor in the endoplasmic reticulum (ER) and delivered via the Golgi complex to immature insulin secretory granules (SGs), also known as (large) dense core vesicles. Activation of prohormone convertases upon acidification of the immature SG lumen drives the conversion of proinsulin into insulin, and thereby SG maturation. Mature insulin SGs are then stored in the cytoplasm until hyperglycemia triggers their exocytosis for insulin release.

Interestingly, insulin SGs are not equally competent for exocytosis, but are categorized into functionally distinct pools. In particular, two factors determine the release probability of SGs. SGs close to the plasma membrane are considered more likely to undergo exocytosis (Rorsma and Renström, 2003; Shibasaki et al., 2007). Equally important but less studied is the age of SGs. Newly-synthesized insulin SGs have a greater propensity for exocytosis compared with their older counterparts (Schatz et al., 1975; Gold et al., 1982; Halban, 1982; Ivanova et al., 2013) - a feature β-cells share with many other cell types showing regulated exocytosis (Kopin et al., 1968; Collier, 1969; Besson et al., 1969; Molenaar et al., 1973; MacGregor et al., 1975; Piercy and Shin, 1981). Aging of SGs correlates with a change of their characteristics: at least in the case of insulin SGs of rat insulinoma INS-1 cells, younger SGs are more mobile and acidic relative to aged SGs (Hoboth et al., 2015; Neukam et al., 2017) - two properties which might contribute to their preferential release. At the same time, older SGs are increasingly removed by autophagy/crinophagy (Marsh et al., 2007; Müller et al., 2017). Proteins or lipids involved in this age-specific behavior, however, are not yet known.

Progress in the molecular characterization of age-distinct SG pools has been hampered by the lack of protocols for their purification. Previous approaches to isolate insulin SGs for proteomic analysis have led to the identification of 51-140 SG proteins, including many established cargoes of these organelles and several new candidates (Brunner et al., 2007; Hickey et al., 2009; Schvartz et al., 2012). However, technical limitations could not prevent the co-enrichment of non-SG proteins, while several well-known SG transmembrane proteins and lumenal cargoes were not detected (Suckale and Solimena, 2010). Typical enrichment protocols for organelles are prone to cross-contamination, which reduces the accuracy of downstream proteomic and lipidomic analyses. The most widely applied method for organelle purification is subcellular fractionation, either by differential or gradient centrifugation. Both procedures, however, cannot avoid the co-enrichment of other cellular compartments, especially vesicular organelles with similar physical properties, such as lysosomes, synaptic-like microvesicles or endosomes. A notable exception has been the purification to homogeneity from brain synaptosomes of neuronal synaptic vesicles, which has been possible largely because of the very high abundance of these organelles (Huttner et al, 1983; Takamori et al, 2006). An alternative approach is the immunoisolation of organelles, with antibodies recognizing a specific epitope in the cytoplasmic domain of a transmembrane bait protein, followed by the binding of the antibody to an affinity matrix (Thomas-Reetz, et al., 1993; Walch-Solimena et al., 1993; Klemm et al., 2009). Since copies of post-Golgi vesicle transmembrane proteins are also in transit through the ER and the Golgi complex at the time of cell lysis, contamination by the latter organelles is nonetheless possible.

Here we report an immuno-based approach for the purification of insulin SGs of distinct age. Our protocol takes advantage of the specificity of immunopurification and combines it with pulse-chase labeling to restrict the antigen to post-Golgi organelles using a cleavable CLIP tag. We show that our approach dramatically reduces background, giving access to highly-purified SGs that can be eluted as intact organelles. Notably, this approach is further suitable for the isolation of age-distinct insulin SG pools. We analyzed the lipidomic profile of young and old SGs from INS-1 cells and found that the ratio of phosphatidylcholine (PC) to phosphatidylethanolamine (PE) changes during SG aging. These changes imply greater membrane fluidity of membranes from aged SGs *in vitro*. Additionally, the proteomic profiles of age-distinct SGs suggest enrichment of the GTPase RAB3a, as well as the motor protein dynein on young SGs.

## Results

### Strategy

Immunopurification protocols provide high specificity for the respective antigen, but often suffer from contamination by pull-down of other compartments and unspecific binding to magnetic beads. To address those shortcomings, we fused the CLIP tag to the cytoplasmic C-terminus of Ptprn2/phogrin, an intrinsic membrane protein of the SGs (Fig. 1A; Wasmeier and Hutton, 1996). CLIP is a self-labeling protein tag which can covalently bind to a cell-permeable CLIP-substrate (such as BC-TMR or BC-Fluorescein; Gautier et al., 2008) in a pulse-chase manner, thereby ensuring that only newly synthesized phogrin-CLIP is labeled. Cells are then incubated until labeled phogrin-CLIP exits the Golgi complex and it is sorted into immature SGs (Fig. 1B), which then typically evolve in mature SGs within 2 hours (Davidson et al., 1988). Varying the time interval between the labeling of newly synthesized phogrin-CLIP and cell lysis, it is therefore possible to obtain extracts in which only a time-resolved pool of SGs contains the labeled phogrin-CLIP bait.

**Fig. 1.**
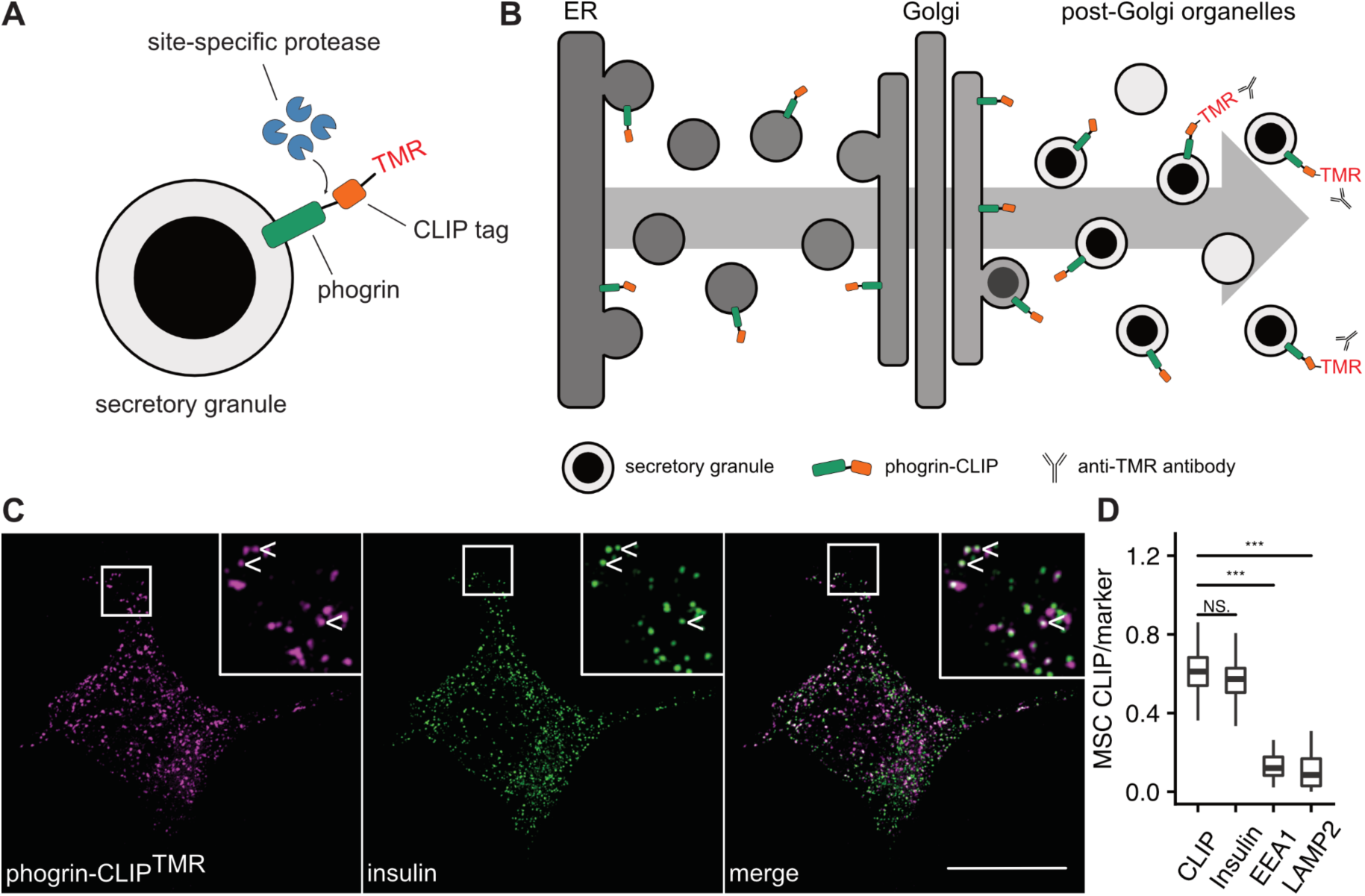
Generation and subcellular localization of phogrin-CLIP. (**A**) Scheme illustrating the construct used to purify insulin SGs. A linker containing a cleavage site for the human rhinovirus (HRV) 3C protease connects the cytoplasmic C-terminus of phogrin with a CLIP tag. (**B**) Phogrin-CLIP enters the secretory pathway and is specifically sorted into SGs at the trans-Golgi network. Labeling with TMR can be timed to label only post-Golgi SGs. Immunopurification is performed with a specific antibody against the substrates (e.g. TMR). (**C**) Representative SIM image of an INS-1 cell stably expressing phogrin-CLIP labeled with CLIP-Cell TMR-Star and co-stained for insulin. (**D**) Quantification of co-localization of phogrin-CLIP labeled with TMR and various markers. Anti-CLIP antibody served as positive control. Co-localization was calculated using the Manders split coefficient (MSC). Data are represented as boxplots with median ± SD. Statistical analysis corresponds to t-test with p-values equaling ***< 0.001, NS. = not significant. Cells from three independent experiments (n = 24-32) were analyzed. Scale bar = 10 µm.

Using an antibody against the CLIP substrate, such as the fluorescent dye tetramethylrhodamine (TMR) or fluorescein (Fluo), rather than against the bait protein allows for both the purification of age-distinct SGs and the exclusion of contaminants from pre-SG compartments along the secretory pathway. To reduce background by non-specific binding to the magnetic beads, we further included a human rhinovirus (HRV) 3C protease cleavage site between the CLIP tag and the cytoplasmic domain of phogrin enabling the specific elution of the immunoisolated organelles from the beads (Fig. 1A). We chose phogrin as a bait protein specific to SGs, because unlike its paralogue PTPRN/islet cell autoantigen 512 (ICA512 -also known as IA-2; Ort et al., 2001; Trajkovski et al., 2004), it is not susceptible to proteolytic cleavage by calpain upon its transient insertion to the plasma membrane (Taraska et al., 2003), and thereby potentially depleted in older SGs following repeated rounds of kiss-and-run or partial exocytosis. Hence, phogrin is an optimal candidate for the cytoplasmic addition of a CLIP tag at its C-terminus. Since phogrin has been proposed to have phosphatidylinositol phosphatase activity (Caromile et al., 2010), we further replaced its catalytic cysteine 931 with a serine (Caromile et al., 2010) to avoid interferences by overexpression of our reporter. Addition of the CLIP tag not only allows for the age-distinct purification of SGs, but also limits the antigen (the CLIP label) to post-Golgi compartments.

### Microscopic and biochemical characterization of phogrin-CLIP

First, we tested the subcellular localization of phogrin-CLIP C931S (henceforth termed phogrin-CLIP) to ensure its efficient and specific targeting to insulin SGs. Super-resolution microscopy revealed that phogrin-CLIP covalently labeled with the dye BC-TMR was found in punctate structures which almost exclusively co-localized with endogenous insulin (Fig. 1C-D). We did not detect significant co-localization with early endosomal and lysosomal markers Eea1 and Lamp2, respectively (Fig. 1D, S1B-C). Co-labeling for CLIP with an antibody served as a positive control (Fig. 1D, S1A) and revealed that not all fusion proteins were positive for TMR, i.e. that washes after the labeling pulse were sufficient to prevent continuous labeling (Fig. S1A).

We also analyzed some of the properties of a stable phogrin-CLIP INS-1 cell line. As expected, we could not detect the CLIP signal in non-transfected control samples, but only in the stable cell line (Fig. S2A). Addition of the ∼20 kDa CLIP tag retarded the electrophoretic mobility of prophogrin by SDS-PAGE to an apparent molecular weight of ∼120 kDa and that of the converted, mature phogrin to ∼80 kDa (Fig. S2A). Probing for untagged phogrin, we found that the stably expressing phogrin-CLIP INS-1 cells retained the expression of endogenous phogrin similar to control cells, as shown by real-time PCR and immunoblot (Fig. S2A-B). Since phogrin-CLIP was expressed more than endogenous phogrin, we tested whether its overexpression affected the levels of other insulin SG components and insulin secretion. Compared to non-transfected INS-1 cells, phogrin-CLIP INS-1 cells did not display obvious changes in the content of chromogranin A (CHGA), prohormone convertase 2 (PC2) or carboxypeptidase E (CPE), as shown by immunoblot (Fig. S2C). However, stable overexpression of phogrin-CLIP correlated with increased glucose-stimulated insulin secretion (GSIS), whereas insulin biosynthesis was unaffected (GSTI, Fig. S2D-E).

We further assessed the N-glycosylation status of phogrin-CLIP to verify its maturation and trafficking. Rat and mouse phogrin are predicted to be N-glycosylated at N553 and N550, respectively. Digestion with endoglycosidase H (EndoH), which only cleaves the N-glycan chain prior to its modification in the cis-Golgi, did not change the electrophoretic mobility of (pro)phogrin-CLIP, but affected an intermediate species (Fig. S2F). In contrast, digestion with PNGaseF, which removes all N-glycans, increased the electrophoretic mobility of all species, consistent with being originally N-glycosylated (Fig. S2F). Hence, mature phogrin-CLIP and prophogrin-CLIP, but not an intermediate species, were insensitive to treatment by EndoH (Fig. S2F). These data indicate that phogrin-CLIP progresses as expected along the secretory pathway.

Next, we transfected the stable phogrin-CLIP line with our previously described human insulin-SNAP reporter for SG aging (Ins-SNAP, Ivanova et al., 2013) to test whether the two reporters have similar localization and SG aging kinetics. When the CLIP and SNAP tags were simultaneously labeled with orthogonal dyes BC-TMR and SNAP-Cell^®^ 430, respectively, for a SG age of 2-4 hours we could detect a high level of correlation (Fig. S3A-C). However, the signals of 20-22 hour-old insulin-SNAP^430^ and 2-4 hour-old phogrin-CLIP^TMR^ were clearly distinct and not significantly correlated (Fig. S3B-C).

Taken together, we conclude that phogrin-CLIP is transferred efficiently to the Golgi and post-Golgi compartments and that its stable overexpression does not significantly affect insulin SG stores or secretion. It also has similar kinetics to Ins-SNAP and can therefore serve as a reliable bait for the immunoisolation of age-distinct insulin SG pools.

### Immunopurification of phogrin-CLIP and SGs

Despite the use of specific antibodies, immunoisolation protocols often suffer from non-specific co-enrichment. This is due to non-specific binding of the antibodies, as well as unintended binding of proteins to the solid phase, such as magnetic beads, during the isolation procedure. To reduce the latter, we incorporated a site-specific protease cleavage site into phogrin-CLIP (Fig. 1A), which allows for the selective elution of the SGs from the beads. Using silver staining after SDS-PAGE, we assessed the degree of background of immunoisolated labeled and unlabeled samples. As expected, the input and flow-through in both TMR positive and negative samples appeared similar (Fig. 2A) and showed a plethora of bands. Similarly, beads from both samples showed distinct protein bands, which, however, were slightly more abundant in the TMR positive samples. The eluates, released from the beads by addition of the protease, were instead remarkably different. Apart from a distinct band of ∼25 kDa, representing the added protease, no protein was visible in the eluate of the beads incubated with the unlabeled phogrin-CLIP control sample. In contrast, the eluate of the beads incubated with the TMR labeled phogrin-CLIP sample showed a multitude of protein bands (Fig. 2A).

**Fig. 2.**
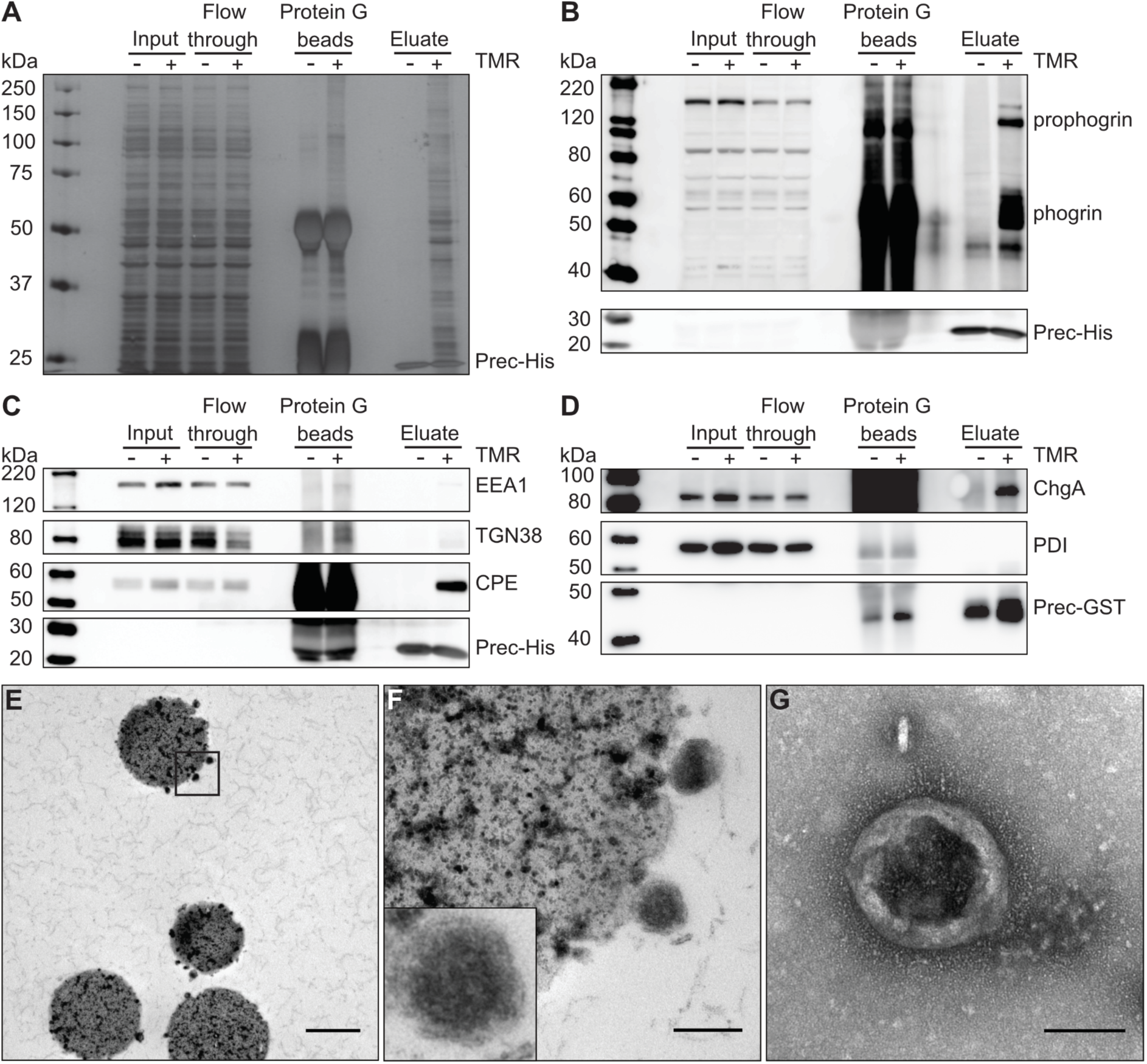
Characterization of SG enrichment purity. (**A**) Silver staining of material enriched from cells labeled with or without CLIP-Cell TMR-Star. Input (10 µg), flow-through (10 µg) and Protein G bead fractions were loaded for comparison. The added protease is visible at ∼25 kDa. (**B**) Material from SG purification was loaded and probed for phogrin by Western Blot. (**C**) SGs were purified and analyzed by Western Blot for the organelle markers Eaa1, Tgn38 and Cpe or (**D**) ChgA and Pdi. The His- or GST-tagged HRV 3C protease (Prec-His or -GST) in the eluate served as loading control. (E-G) Ultrastructural analysis of purified insulin SGs by electron microscopy. (**E**) Micrograph of a bead sample prior to the addition of the HRV 3C protease embedded in epon. (**F**) Micrograph of inset from (E). (**G**) Negative staining of eluted material from specific samples as shown in (E, F). Scale bar = 1 µm (E), 200 nm (F), 100 nm (G).

To verify the selective immunoisolation and elution of SGs, we immunoblotted control and labeled fractions for markers of distinct intracellular compartments. In contrast to the control sample, the eluate from the TMR labeled phogrin-CLIP sample was enriched for several granule cargoes, including CPE, PC2, CHGA, ICA512 and phogrin (Fig. 2B-D, S4A-C). It was negative instead for Golgi markers TGN38 and GM130, ER markers CANX and PDI (Fig. 2C-D, S4A-B), as well as the early endosome marker EEA1 (Fig. 2C). Interestingly, we could also detect the immature SG marker islet cell autoantigen 69 (ICA69, Spitzenberger et al., 2003; Buffa et al., 2008, Fig. S4A). As expected from the silver staining, none of the mentioned markers were detected in the eluate from the control beads. We could, however, detect the lysosomal marker LAMP2 and the synaptic-like microvesicle marker synaptophysin 1 (SYP1, Fig. S4D). To limit the contamination by these organelles, we included an immunodepletion step with antibodies targeting the cytoplasmic tails of either LAMP2 alone or both LAMP2 and SYP1. Interestingly, these two proteins could be depleted by the immunoisolation with the anti-LAMP2 antibody alone, suggesting that both markers resided in a common organelle (Fig. S4E).

Finally, we evaluated the ultrastructural integrity of the purified SGs by transmission electron microscopy. To this aim either beads of TMR-labeled samples were embedded in epon for imaging of ultrathin sections (Fig. 2E-F), or eluted SGs were directly spotted on grids for negative staining (Fig. 2G). Electron micrographs confirmed the purity and integrity of the immunoisolated SGs, with no obvious contamination from other organelles. Epoxy-embedded SGs bound to beads contained the characteristic dense core of insulin granules surrounded by a lipid bilayer (Fig. 2F). In summary, these data indicate that our protocol allows for the highly specific and background-free enrichment of insulin SGs from INS-1 cells using a cytoplasmic CLIP tag.

### Lipidomic analysis of isolated insulin SGs

Having established that our protocol is suitable for the purification of SGs with high purity, we isolated SGs for shotgun lipid mass spectrometry (MS). For isolation of SGs without age-bias, we labeled the cells without addition of a blocking substrate, thereby labeling all pre-existing SGs (termed ‘mixed’). To investigate potential differences in lipid profiles of young and old SGs, we first labeled phogrin-CLIP with an additional blocking step to saturate all existing CLIP (Fig. 1B). We then labeled newly-synthesized phogrin-CLIP with BC-Fluo and incubated the cells until labeled phogrin-CLIP reached the desired age: 1-4 h for young SGs and 16.5-22 h for old SGs.

Given the overall low amount of sample that was available for MS, we were able to identify with high certainty ∼100 different lipid species (Tab. S1, S2). These likely represent the most abundant species. When analyzing the data by principal component analysis, we found that samples from young SGs clustered well together and separately from the similarly clustered old SGs, whereas mixed samples were scattered in between (Fig. 3A).

**Fig. 3.**
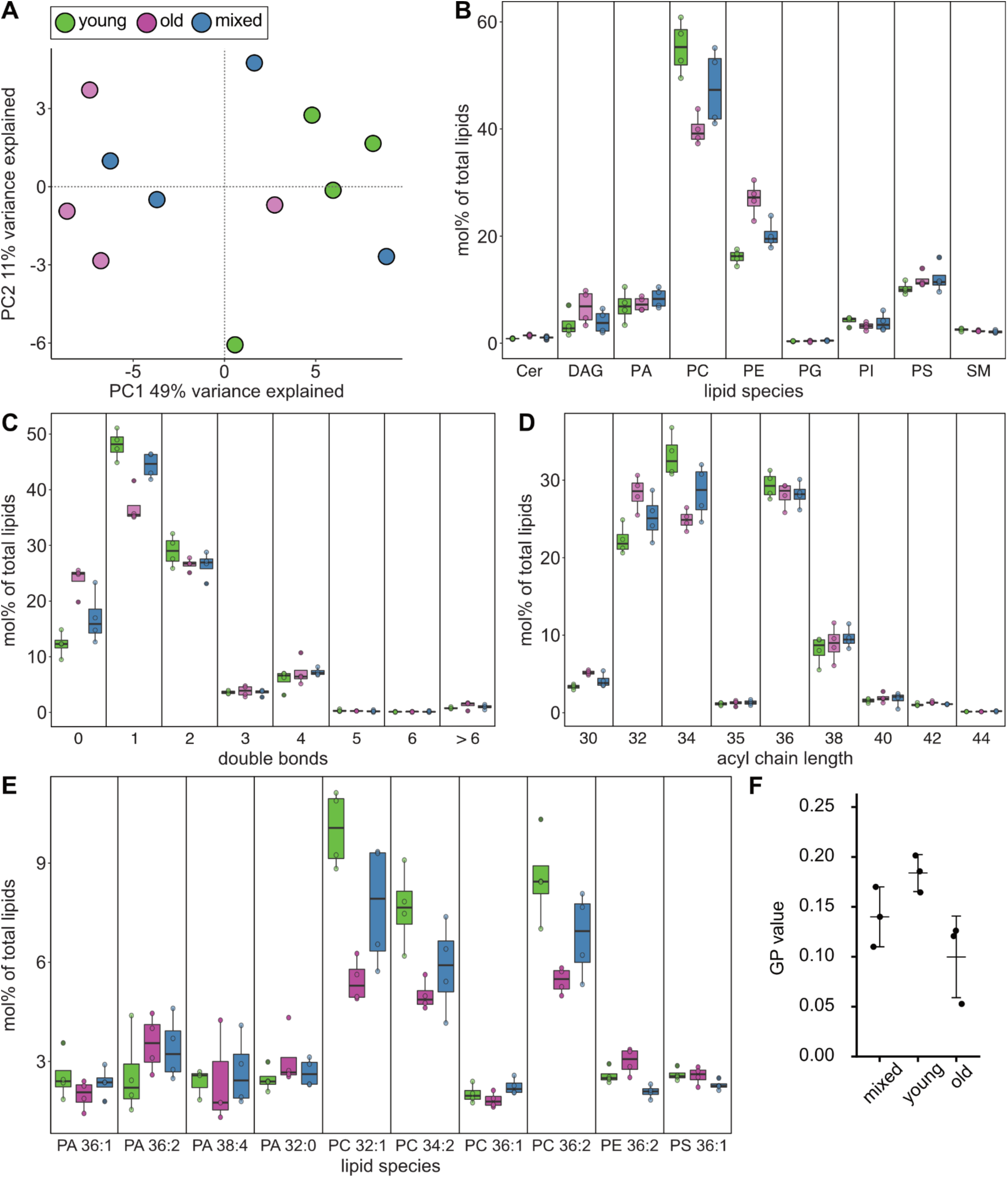
Lipid profile of purified young and old SGs. (**A**) Principal component analysis of individual samples of young (green), old (magenta) and mixed (blue) SGs. (**B**) Lipid classes detected in preparations of young (green), old (magenta) and mixed (blue) SGs. Boxplots are shown for ceramide (Cer), diacylglycerol (DAG), phosphatidic acid (PA), phosphatidylcholine (PC), phosphatidylethanolamine (PE), phosphatidylglycerol (PG), phosphatidylinositol (PI), phosphatidylserine (PS) and sphingomyelin (SM). (**C**) Distribution of the number of double bonds identified in lipids from young (green), old (magenta) and mixed (blue) SGs. (**D**) Distribution of acyl chain length identified in lipids from young (green), old (magenta) and mixed (blue) SGs. (**F**) General polarization (GP) values of membranes from mixed, young and old SGs. Measurements from three independent SG preparations.

Analysis of the lipid profiles of purified insulin SGs revealed a strong contribution of phosphatidylcholine (PC), phosphatidylethanolamine (PE) and phosphatidylserine (PS, Fig. 3B). We further detected considerable amounts of diacylglycerol (DAG) and phosphatidic acid (PA). Minor fractions of phosphatidylinositol (PI) and sphingomyelin (SM), as well as negligible amounts of ceramide (Cer) and phosphatidylglycerol (PG) were present (Fig. 3B). Interestingly, we could measure compositional differences between young and old SG fractions, with a higher fraction of PC in young and PE in old SGs (Fig. 3B). Most of the lipids were unsaturated with either one double-bond (46.7 ± 2.67% and 36.53 ± 2.90% for young and old SGs, respectively) or two double-bonds (28.53 ± 2.81% and 26.14 ± 1.25% for young and old SGs, respectively, Fig. 3C). Only 12.03 ± 2.12% and 23.48 ± 2.10% of the lipids were fully saturated in young and old SGs, respectively (Fig. 3C). The overall length distribution appeared to favor chain lengths of 32, 34 and 36 carbons and a smaller fraction of 38 (Fig. 3D). Taken together, old SGs contained a higher fraction of lipids with higher saturation and shorter acyl chain length (Fig. 3C-D). We analyzed the ten most abundant lipid species for potential changes between young and old SGs. In old SGs, PE 32:0 was more abundant, whereas young SGs contained more PC 32:1, 34:1, 34:2 and 36:2 (Fig. 3E).

To evaluate whether the enrichment of PE in old and PC in young SGs has an influence on membrane properties, we measured the membrane fluidity. To do so, we prepared liposomes from lipid extracts of isolated SGs in which the fluorescent c-Laurdan probe was introduced to measure the General Polarization (GP, Kaiser et al., 2009) index. We found that liposomes formed from lipids of young SGs were more rigid than those of old SGs, as evidenced by their higher GP values (Fig. 3F). Notably, the composition of mixed SGs had intermediate lipidomic (Fig. 3B-E) and GP values (Fig. 3F), presumably reflecting a true mixture of young and old SGs.

In summary, the lipidomes of young and old SGs revealed a change of the PC/PE ratio during the granule aging process. This difference is sufficient to entail an *in vitro* change of the biophysical properties of their membranes.

### Proteomic analysis of isolated insulin SGs

To characterize the protein composition of our purified SGs, we labeled SGs as described above and examined the eluted material by MS. First, we purified total SGs without age-bias and analyzed their proteome in comparison to control samples without labeling. Similar to the analyses by silver gel and Western Blot, we found that the unlabeled samples contained significantly less proteins than the labeled fractions (Fig. 4A-B). The 299 proteins identified in unlabeled fractions were amongst the 1010 proteins present in labeled fractions and no protein exclusive to unlabeled samples was identified (Fig. 4A-B). Significantly enriched were 8 proteins in unlabeled and 875 in labeled fractions (Fig. 4A, Tab. S4). In the labeled fractions, we could identify most known SG proteins, such as CPE, PTPRN/ICA512, PTPRN2/phogrin, INS1/2, PCSK1/2, IAPP, VAMP2 and subunits of the v-ATPase (Fig. 4A). However, despite pre-clearing, lysosomal contaminants, such as LAMP1/2 or cathepsins A/B could still be detected (Fig. S6A). Additionally, markers of endosomes (RAB7a, RAB11) and mitochondria (VDAC2, ATP5a1) were also present.

**Fig. 4.**
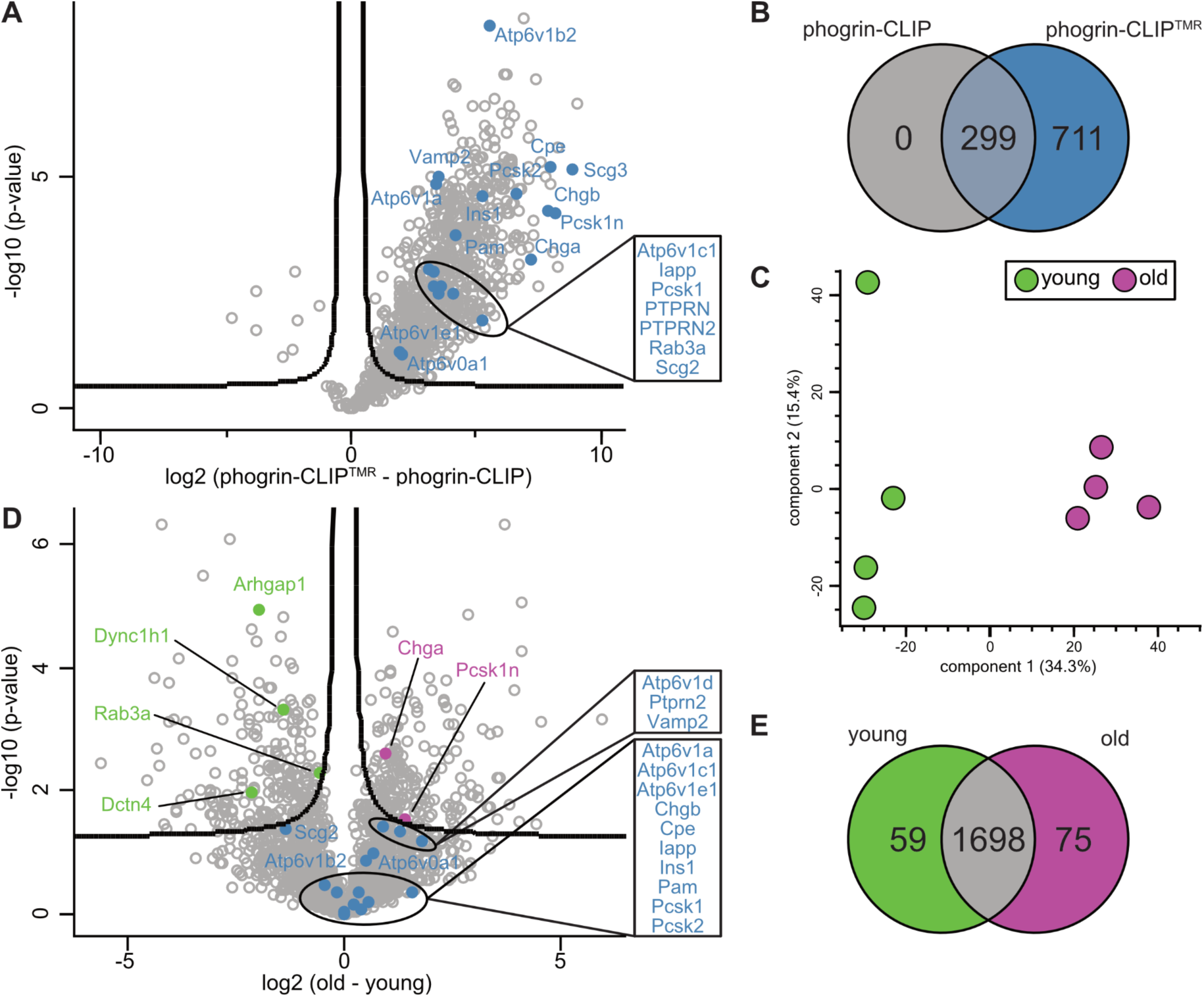
Proteomic characterization of purified SGs. (**A**) Volcano plot showing proteins significantly enriched in unlabeled (left) and labeled (right) samples, with FDR 0.05 and S0 0.1. Data represent median values of four independent experiments. (**B**) Venn diagram showing the amount of shared or exclusive proteins as identified in (A). (**C**) Principal component analysis of the individual proteomic profiles of young (green) and old (magenta) SGs. (**D**) Volcano plot showing proteins significantly enriched in preparations of young (left, green) or old (right, magenta) SGs, with FDR 0.05 and S0 0.1. SG proteins not significantly changed are shown in blue. Data represent median values of four independent experiments. (**E**) Venn diagram showing the amount of shared or exclusive proteins as identified in (D).

Similar to the lipidomic profiling, we analyzed the proteomes of young and old SGs. Unbiased principal component analysis indicates that also on the proteomic level young and old SGs formed well-defined clusters clearly distinct from each other (Fig. 4C). Importantly, the majority of known SG proteins, such as INS1/2, IAPP, PCSK1/2 and PAM, were not enriched in either of the two pools, suggesting that their preparations yielded similar amounts of SGs (Fig. 4D). However, the small GTPase RAB3a was enriched in young SGs, whereas CHGA and PCSK1n/ProSAAS were significantly enriched in old SGs (Fig. 4D).

When comparing the overall similarities between the samples, we found that 1698 proteins were shared amongst the eluates of young and old SGs, whereas 59 and 75 proteins were exclusive for the eluates of young or old SGs, respectively (Fig. 4E). Significantly enriched were 222 proteins in fractions of young and 188 in fractions of old SGs (Tab. S6). Amongst the shared proteins, cytoplasmic dynein 1 heavy chain 1 (DYNC1h1), as well as its adaptor dynactin subunit 4 (DCTN4) were enriched on young SGs (Fig. 4D, Tab. S6). The Rho GTPase-activating protein 1 (ARHGAP1), another cytosolic protein potentially involved in SG exocytosis, was also significantly enriched on young SGs. Importantly, the lysosomal markers LAMP1/2, cathepsins A/B and the endosomal markers RAB7a and RAB11 were not enriched in either fraction, pointing to comparable contamination of the two age-distinct SG pools (Fig. S5B).

We further examined proteins exclusive to eluates of either young or old SGs (Fig. S5C, Tab. S7). Proteins present only in eluates of young SGs comprised mostly ribosomal (e.g. RPL4) or RNA binding proteins (e.g. HNRNPC, EIF2s2), whereas in eluates of old SGs mitochondrial proteins (e.g. ALDH1l2, GRPEL2, CMC1) were present.

Taken together, these data indicate that most SG proteins, in particular lumenal and transmembrane proteins, remain constant during the aging process. However, cytosolic proteins, such as motor proteins or small GTPases, may dynamically interact with SGs of distinct age.

## Discussion

### Antigen restriction allows for isolation of age-distinct secretory granules

Despite sophisticated combinations of density centrifugation (Brunner et al., 2007) and additional combination with SILAC (Schvartz et al., 2012), isolation of insulin granules could not efficiently avoid contamination by non-granular organelles with similar physical properties. Alternatively, organellar specificity can be achieved using antibodies targeting transmembrane proteins such as the SNARE protein VAMP2 (Hickey et al., 2009). Yet, also this approach suffers from a major drawback: the bait protein may not be entirely specific for an organelle, as is the case for VAMP2, which is also present on synaptic-like microvesicles and recycles through endosomes multiple times. Further, SG proteins naturally travel along the secretory pathway. Hence, at the time of cell disruption, any such cargo will inevitably also be found on pre-granular organelles such as the Golgi or the ER, resulting in their co-enrichment.

To limit these contaminations, we took advantage of the specific sorting of phogrin to SGs and the attached CLIP tag for purification. Rather than using antibodies against CLIP, we use its substrate as antigen. With pulse-chase labeling of the CLIP tag with TMR- or Fluorescein-conjugated substrates, and a preceding blocking step with the unlabeled substrate, only newly-synthesized phogrin-CLIP is marked. Given enough time for the fusion protein to exit the TGN for sorting on nascent SGs, the antigen is restricted to post-Golgi organelles. This avoids the co-purification of notorious contaminants, such as the Golgi and ER, which were largely absent in our purified material. Culturing the cells at glucose levels not eliciting secretion further limited the number of SGs undergoing exocytosis and therefore the amount of phogrin-CLIP^TMR^ transiently resident in the plasma membrane and endosomal compartments (Vo et al., 2004; Wasmeier et al., 2005). Remaining contaminants were lysosomes, which are responsible for the intracellular disposal of SGs either through autophagy or crinophagy (Orci et al., 1984; Marsh et al., 2007; Riahi et al., 2016; Müller et al., 2017). These could be largely immunodepleted with antibodies directed against the cytosolic tail of the specific lysosomal membrane protein LAMP2. As a result, we were able to isolate intact SGs with a high degree of purity. In addition, our approach allows for the age-specific isolation of SGs, as the time between labeling and harvesting of cell lysates can be varied to the desired granular age.

To our knowledge, this is the first time that age-distinct pools of a given cellular organelle could be isolated for biochemical analysis. The secretory granule is the optimal and perhaps only organelle lending itself to this opportunity, being largely a stationary compartment and presumably undergoing only limited remodeling after biogenesis at the TGN. Additionally, its lifetime of several days (Halban and Wollheim, 1980; Müller et al., 2017) and presence in many hundreds of copies in each peptide-hormone/neuropeptide secreting cell are beneficial for its time-resolved purification. This possibility seems instead precluded, and conceivably less biologically insightful, for other organelles such as mitochondria, peroxisome and compartments of the secretory or endolysosomal pathway, whose membrane protein composition undergoes a rapid turnover, remodeling by fusion and fission, and/or which have a very short half-life.

As phogrin is widely expressed in peptide-hormone and neuropeptide secreting cells, our protocol could be easily applicable to other cell types and lines, hence helping to better understand the mechanisms of their regulated secretion.

### Lipidomic analyses of age-distinct secretory granules

To the best of our knowledge, only a single study addressed the lipidomic composition of insulin SGs (MacDonald et al., 2015). In this approach, lipids were separated by thin layer chromatography and the fatty acid composition of each lipid class was analyzed by gas chromatography mass spectrometry (GC-MS). Here, we applied shotgun MS to whole lipid extracts together with internal standards for each lipid class. We identified one lipid class that was not observed by the previous study: PA. It is a low abundant phospholipid in cell membranes, which acts on one hand as a building block for all other phospholipids, and on the other hand as an important player in cellular signaling and as a recruitment factor for specific proteins. The levels of the other phospholipids measured here also differed in comparison to the previous study, especially PC 46% vs. 22%, PE 21% vs 17%, PS 12% vs 11%, PI 4% vs. 21% and SM 3% vs. 10%. Such substantial differences can be attributed to the methodology (solid-phase extraction, GC-MS vs. direct infusion) and sample purity (differential centrifugation vs. affinity isolation) or even minor differences, like the serum used for cell culture and other related variables.

Notably, our data are to some extent consistent with the lipid levels found in chromaffin granules (Blaschko, et al., 1967; Winkler et al., 1967; Da Prada et al., 1972; Balzer and Khan, 1975; De Oliveira Filgueiras et al., 1981), where PC was 25-28% (bovine)/37% (rat), PE was 27-36% (bovine)/30% (rat), PS was 7.9-8.9% (bovine)/11% (rat), PI was 1.5-2.9% (bovine), SM was 11-20% (bovine)/9% (rat). Nevertheless, the overall composition is similar to that of the plasma membrane, which seems plausible, considering that the effect of exocytosis on PM lipid homeostasis should be limited (Sezgin et al., 2015).

Comparing the lipid profiles of young and old SGs, we found changes in the most abundant lipids. Young SGs contained higher levels of PC 32:1, 34:1, 34:2 and 36:2, whereas old SGs had higher levels of PE 32:0. These variations were associated with a greater membrane fluidity *in vitro* for liposomes formed from old SGs. Such changes may dynamically regulate the association of cytosolic proteins to the SG membrane, as well as the activity of SG transmembrane proteins, e.g. various channels and transporters.

The observed relative changes in lipid content of the old vs. young SGs were restricted to very few, specific lipid species, indicating a directed process for their exchange/enrichment. An enzymatic change of the lipid headgroups seems unlikely, as the respective acyl chains were not maintained. A plausible explanation is therefore the transfer of particular lipid species at contact sites between SGs and other membrane organelles, as it has been described to occur at physical proximity sites between the PM and ER, endosomes and ER or lysosomes and peroxisomes (reviewed in Eisenberg-Bord et al., 2016; Saheki and De Camilli, 2017). This potential lipid transfer might be of very transient nature and future work will be required to address this possibility.

### Proteomic analyses of age-distinct secretory granules

Various approaches have been used to characterize the insulin SG proteome, including techniques like density centrifugation with (Schvartz et al., 2012) and without SILAC (Brunner et al., 2007) or affinity purification with anti-VAMP2 antibodies (Hickey et al., 2009). All three studies were able to detect prominent SG constituents, such as INS1/2, CPE, PCSK2 or CHGA. However, cargoes like PTPRN, PTPRN2, PCSK1n and CHGB were only detected in some of the studies. We complement these findings with our dataset, which additionally includes the previously missed islet amyloid polypeptide (IAPP).

In total, we identified 875 proteins significantly enriched in SG fraction. As it is unlikely that SGs contain 875 specific proteins, the vast majority of the enriched proteins are presumably non-SG proteins. Two reasons may be responsible for the detection of such a large number of proteins. First, the negative control was virtually devoid of proteins. As a result, even minor contaminating organelles in which phogrin-CLIP^Fluo^ was transiently present during homogenization will appear as significantly enriched. Second, advances in mass spectrometry sensitivity may greatly enhance the detection of proteins in general (Brunner et al., 2021). Presumably a combination of the two factors is responsible for the results presented here. Nevertheless, this sensitivity may be crucial to detect well-known, but previously undetected SG proteins like IAPP.

Whilst we detected several single-pass transmembrane proteins of SGs (for example PTPRN, PTPRN2, PAM), multi-pass transmembrane proteins were missed, including the well-known and disease-relevant SG zinc transporter 8 (SLC30a8, Chimienti et al., 2004). The low abundance of such multi-pass transmembrane proteins may fail to produce enough peptides for their unequivocal measurement by mass spectrometry, hence requiring detection and validation by other means.

Analyzing the proteomic differences between young and old SGs, we identified a number of proteins enriched in either age. As expected, the amounts of most lumenal cargoes (e.g. insulin, convertases, granins) and intrinsic single-pass membrane proteins (e.g. PTPRN, PTPRN2, PAM) of SGs remained unchanged during the aging process. Interestingly, however, cytoplasmic factors associated with membrane trafficking, such as the GTPase RAB3a or the motor protein dynein were enriched on young compared to old SGs. The enrichment of motor proteins is consistent with our observation that in β-cells microtubules are prevalently found beneath the plasma membrane and run mostly parallel to it (Müller et al., 2021), therefore ideally positioned to support the bidirectional movement of cortical young SGs seeking for sites of exocytosis (Hoboth et al., 2015). The enrichment of RAB3a, which plays an important role in exocytosis (Regazzi et al., 1996; Yaekura et al., 2003), is also consistent with this scenario. In fact, we could not detect any secretion-associated cytosolic proteins enriched on old SGs, suggesting that during aging SGs lose competence for interaction with soluble cytosolic factors. The reason for such a “shedding” process is not readily clear but changes in the composition of their lipid bilayer, including the lipid headgroups, could reduce the affinity for peripherally associated membrane proteins. For example, the Rab proteins 1/5/6 were shown to specifically insert into negatively charged membranes with lipid packing defects (Kulakowski et al., 2017). It may therefore be of great interest to investigate whether the membranes of young SGs alone are the preferred substrate for RAB3a. In addition, post-translational modifications of proteins, such as phosphorylation, but also many others, could play a crucial role in attracting effectors and future work should address this possibility. Surprisingly, the SG proteins PCSK1n and CHGA were enriched in eluates of old SGs, while the closely related proteins CHGB or PCSK1/3 and PCSK2 remained similar. Being sorted into the lumen of SGs together with all other cargoes, a change in their content over time is unlikely. The enrichment might therefore simply be due to occurrence in contaminating organelles, such as Golgi vesicles or degradative organelles.

Yet, our data indicates the exclusive enrichment of ribosome/ER-associated proteins in the eluates of young SGs, as well as of Golgi-associated and mitochondrial proteins in those of old SGs. This may reflect a young/old specific background, with a minor pool of young phogrin-CLIP^Fluo^ still being resident in ER membranes at the time of purification. Similarly, more of the old phogrin-CLIP^Fluo^ may be found in degradative compartments, such as multivesicular bodies, also containing Golgi or mitochondrial proteins.

### Limitations of this study and outlook

Isolation of subcellular organelles requires large amounts of starting material, for which immortalized cell lines are usually the preferred source (Brunner et al., 2007; Hickey et al., 2009; Schvartz et al., 2012). Although the general composition of organelles from cell lines likely resembles that of their counterparts in primary cells, qualitative and quantitative differences may exist. Quantitative assessments therefore require accurate validation in primary cells. In particular, to investigate age-related changes of insulin granules in a more native environment, the methodology presented here could therefore be transferred to reporter animal models. Although we developed mouse models for the study of SG aging (Ivanova et al., 2013), the low yield of mouse islets and therefore SGs limits their exploitation for the biochemical analysis presented here. Larger animals, such as transgenic pigs, could in this regard be more suitable donors of large quantities of starting material. In fact, we have recently shown that *in vivo* labeling of Ins-SNAP is feasible in transgenic pigs (Kemter et al., 2021). Generation of a β-cell-restricted phogrin-CLIP transgenic pig may therefore be a source to purify age-distinct insulin SGs from primary β-cells.

Another limitation of this study is the exposure of cells to low glucose to suppress their secretion after SG labeling. Since some cytosolic factors may transiently associate with SG membranes only upon physiological stimuli, additional measures would be required to prevent exocytosis while the cells are being stimulated. This could include pharmacological inhibition of secretion or the generation of secretion-deficient cell lines, although this may also enhance the rate of intracellular SG degradation (Marsh et al., 2007).

This purification methodology may be further exploited for metabolomic analyses, e.g. to investigate neurotransmitters stored in SGs or peptides resulting from the conversion of prohormones too small to be identified by other means. Other applications are possible, including the characterization of post-translational modifications of SG proteins, the structural analysis of purified SGs by cryoelectron microscopy or the study of transmembrane proteins in their native environment.

## Acknowledgments

We thank Bert Nitzsche for advice for SIM imaging and Maria Heier for technical contributions in developing the protocol, Andreas Müller for initial electron microscopy, Carolin Wegbrod and Anke Sönmez for cell culture work. We are also grateful to Bettina Mathes for support with the synthesis of BG-Fluorescein(piv)_2_, Howard Davidson for the gift of anti-phogrin antibodies and Mrs. Katja Pfriem for administrative support. Work in the Solimena and Coskun labs was supported by the German Center for Diabetes Research (DZD e.V.), which is financed by the German Ministry for Education and Research, the German-Israeli Foundation for Scientific Research and Development (GIF; grant I-1429-201.2/2017) and the Innovative Medicines Initiative 2 Joint Undertaking under Grant Agreement 115881 (RHAPSODY), which includes financial contributions from the European Union’s Framework Program Horizon 2020, the European Federation of Pharmacological Industries and Associations (EFPIA), the Swiss State Secretariat for Education, Research, and Innovation under Contract 16.0097. The Electron Microscopy Facility is supported by EFRE (European Fund for Regional Development). M.N. was the recipient of a predoctoral fellowship from the Dresden International Graduate School for Biomedicine and Bioengineering (DIGS-BB).

## Author contributions

Conceptualization, M.N., M.S; Methodology, M.N.; Validation, M.N., A.-D.B. and M.G.; Formal analysis, M.N., M.G., J.V., A.-D.B., P.S. and A.P.; Investigation, M.N., A.-D.B., P.S. K.G., M.G., J.V. and T.K.; Resources, J.B. and K.J.; Writing - Original Draft, M.N.; Writing - Reviewing & Editing, M.N., M.G., Ü.C., M.S.; Visualization, M.N. and A.P.; Supervision, Ü.C., M.M. and M.S.; Funding Acquisition, M.N. and M.S.

## Supplementary figures and tables

**Fig. S1.**
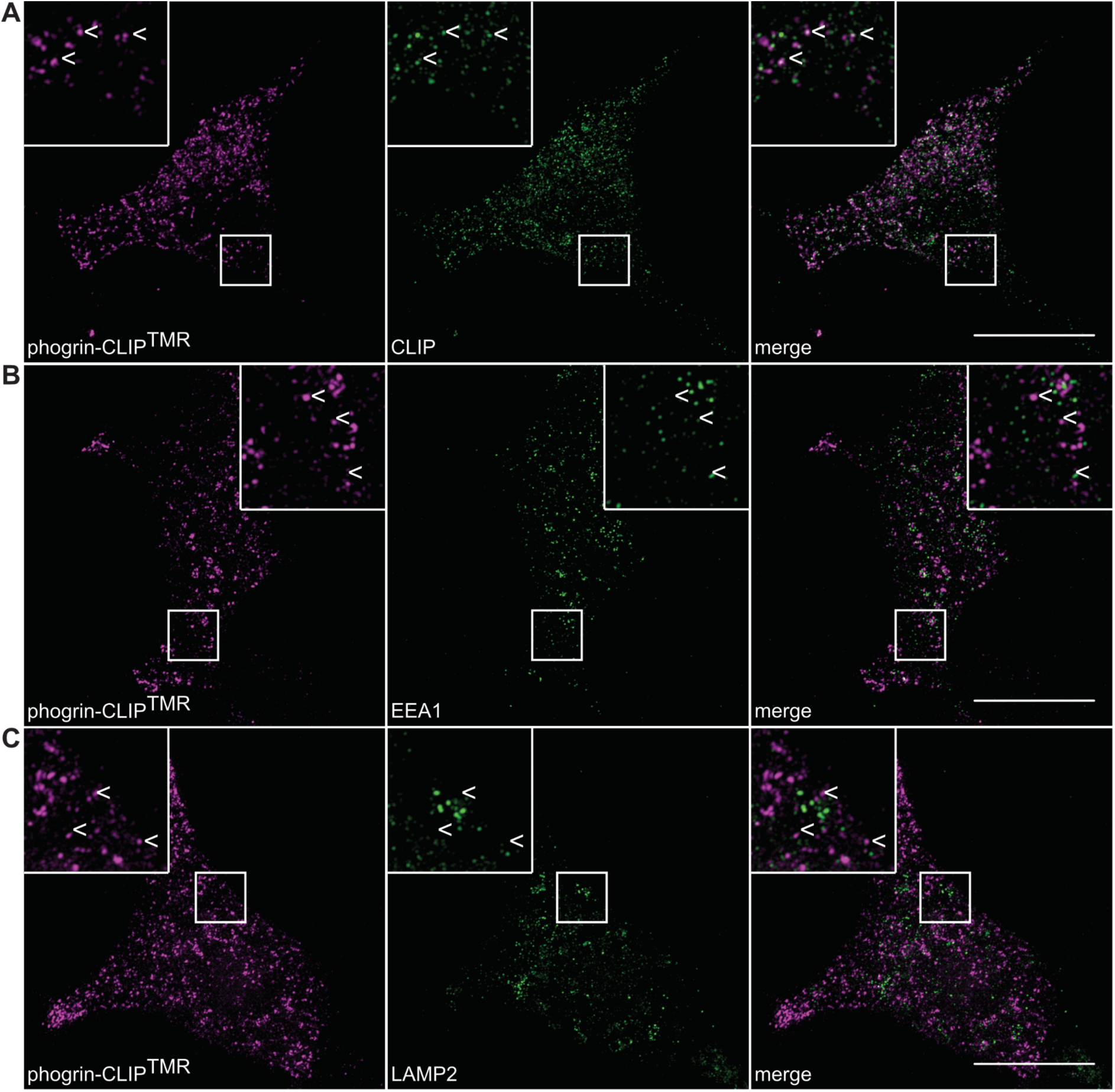
Subcellular localization of phogrin-CLIP. Representative SIM images of INS-1 cells stably expressing phogrin-CLIP labeled with CLIP-TMR and co-stained for CLIP tag (**A**), EEA1 (**B**) and LAMP2 (**C**). Scale bar = 10 µm.

**Fig. S2.**
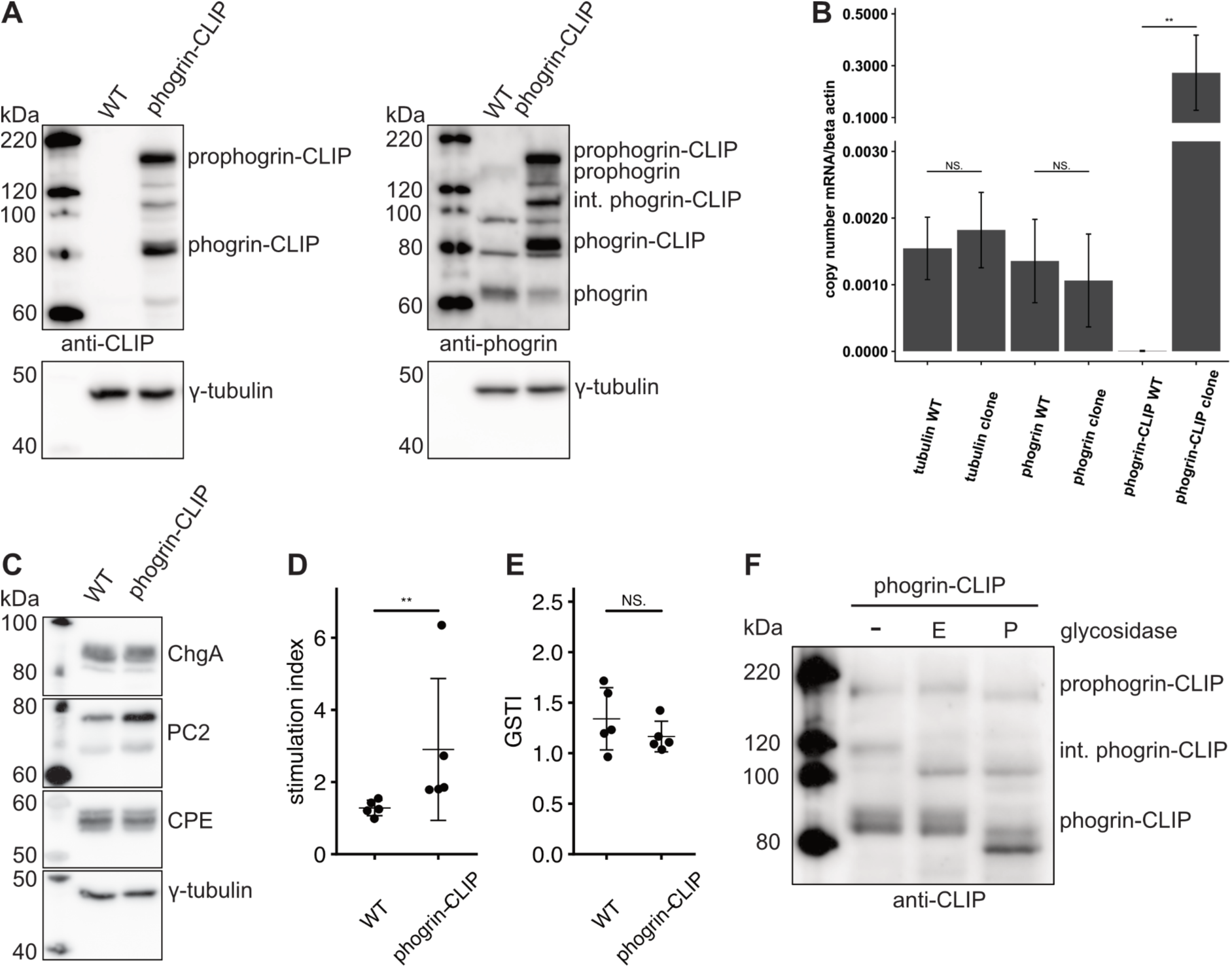
Biochemical characterization of the stably expressing phogrin-CLIP INS-1 clone. (**A**) Cell lysates of non-transfected INS-1 cells and the phogrin-CLIP INS-1 clone probed for CLIP tag (left) or endogenous phogrin (right). (**B**) Real-time PCR of γ-tubulin, endogenous phogrin and overexpressed phogrin-CLIP. The mRNA was normalized to β-actin mRNA. Mean ± SD from 5 independent replicates of non-transfected INS-1 cells and the phogrin-CLIP line. Statistical significance was calculated using the Mann-Whitney-Wilcoxon test with p-values equaling *< 0.05, **< 0.01, NS. = not significant. (**C**) Cell lysates of INS-1 cells and the phogrin-CLIP line were probed for SG markers PC2, CPE and CHGA. Tubulin served as a loading control. (**D, E**) Glucose-stimulated insulin secretion was compared between non-transfected INS-1 cells and the phogrin-CLIP line. The stimulation index and glucose-stimulated total insulin (GSTI) are shown as mean ± SD for 5 independent replicates. Statistical significance was calculated using the Mann-Whitney-Wilcoxon test with p-values equaling *< 0.05, **< 0.01, NS. = not significant. (**F**) Cell lysates of the phogrin-CLIP line were either left untreated (-), treated with endoglycosidase H (E) or PNGaseF (P). Differences in glycosylation were detected by electromobility shift in SDS-PAGE and subsequent probing by Western Blot with antibodies recognizing the CLIP tag.

**Fig. S3.**
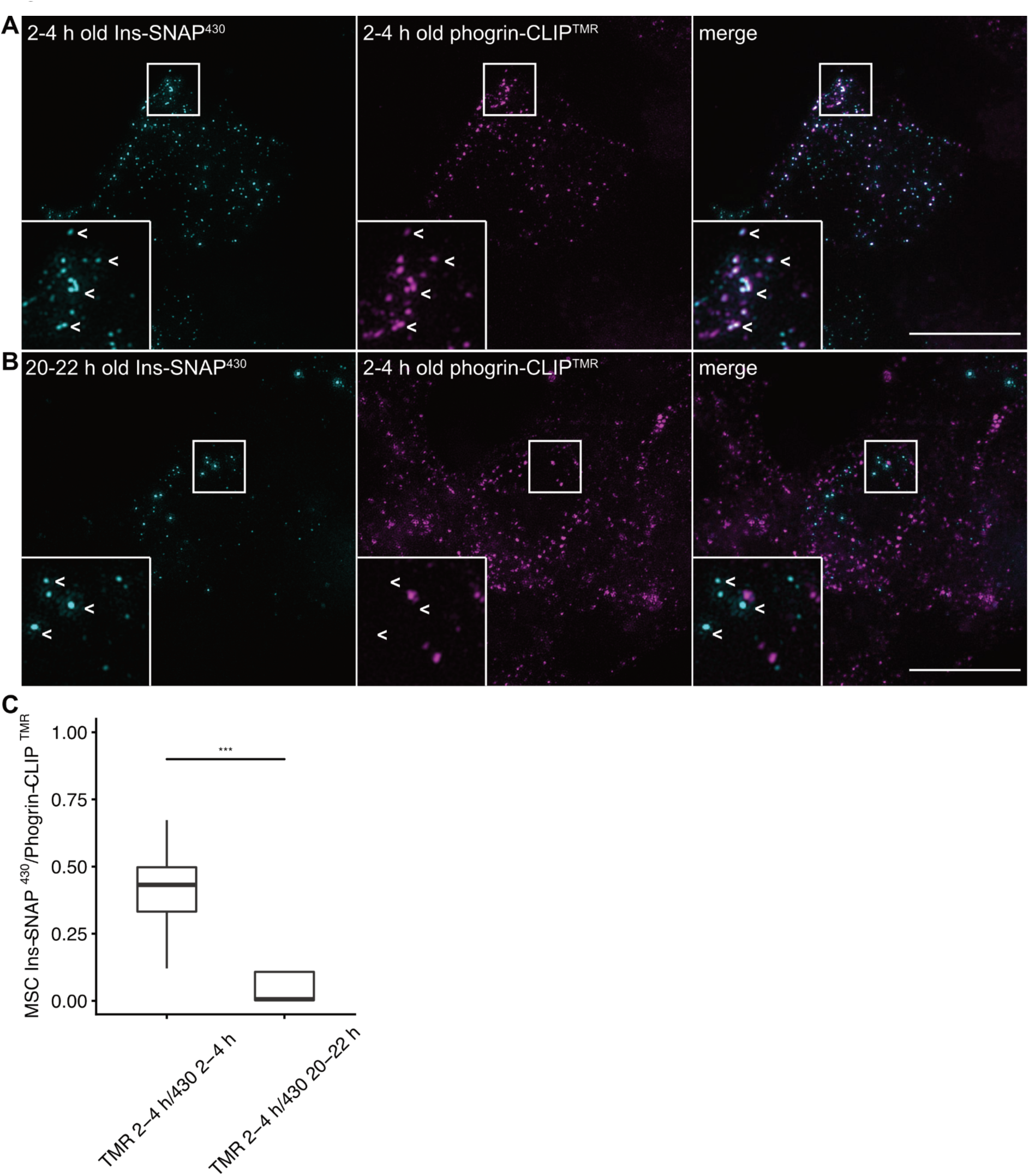
Comparison of aging kinetics of phogrin-CLIP and Ins-SNAP. The phogrin-CLIP line was transfected with Ins-SNAP and labeled with SNAP-Cell^®^ 430 and BC-TMR. (**A**) Representative SIM image when both tags were labeled for an age of 2-4 h or (**B**) 2-4 h for phogrin-CLIP and 20-22 h for Ins-SNAP. (**C**) Quantification of co-localization as calculated by Manders split coefficient (MSC). Data are represented as boxplots with median ± SD. Statistical analysis corresponds to t-test with p-values equaling ***< 0.001, NS. = not significant. Cells from three independent experiments (n = 21-23) were analyzed. Scale bar = 10 µm.

**Fig. S4.**
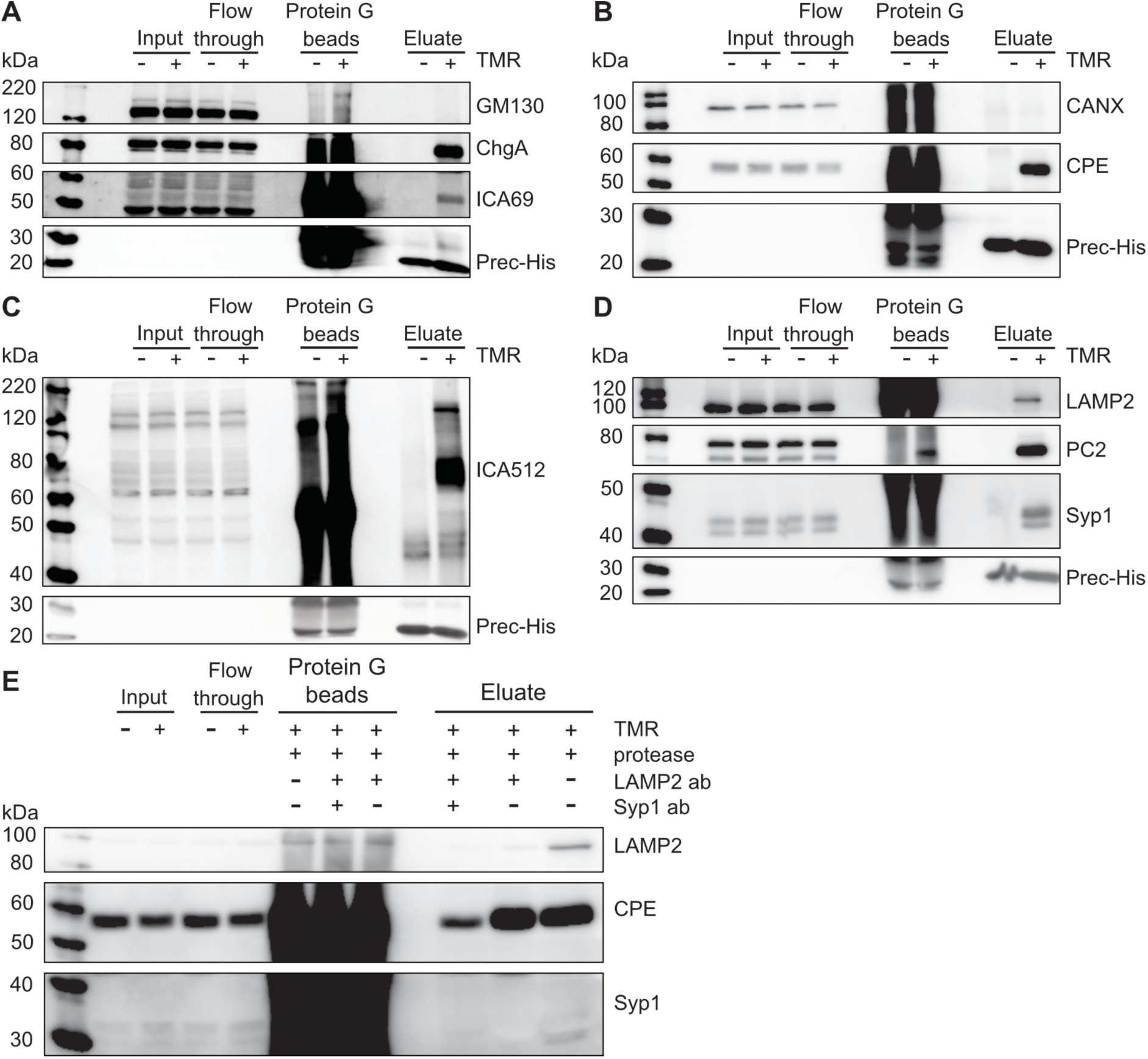
Characterization of enrichment purity. SGs were purified and probed for different organelle markers by Western Blot. Eluates of samples labeled with, or without TMR were probed for SG markers CHGA (**A**), ICA69 (**A**), ICA512 (**C**) and CPE (**B**). Eluates were further probed for the Golgi marker GM130 (**A**) and the ER marker CANX (**C**). His-tagged HRV 3C protease (Prec-His) served as loading control. (**D**) Purified SG lysates were probed for the lysosome marker LAMP2, the SG marker PC2 and the SLMV marker SYP1 by Western Blot. (**E**) SG purification with an additional second immunodepletion step with antibodies either against LAMP2 alone or in combination with SYP1.

**Fig. S5.**
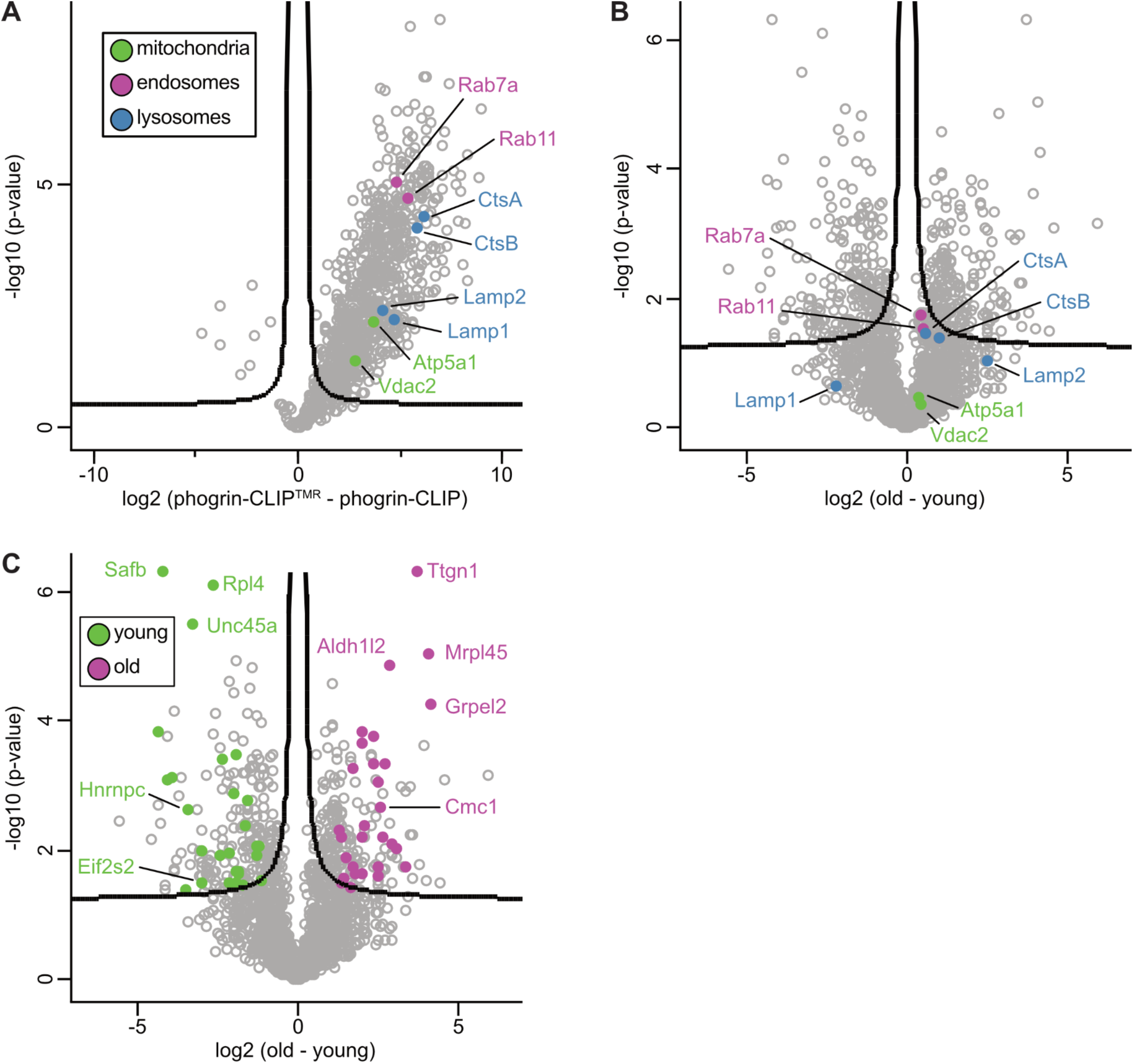
Proteomic characterization of purified SGs. (**A**) Volcano plot showing mitochondrial (green), endosomal (magenta) and lysosomal (blue) proteins significantly enriched in labeled samples. (**B**) Volcano plot showing the same proteins as in (**A**) in samples of young and old SGs. (**C**) Volcano plot showing proteins exclusive for samples of young (green) and old (magenta) SGs. All data represent median values of four independent experiments, with FDR 0.05 and S0 0.1.

**Tab. S1 Lipids identified by MS from eluates of SGs.** Cells were labeled with Fluo for mixed, young or old SGs and the eluates measured by MS. The data represent four biological replicates and were filtered for a coverage of 75%. Data are in pmol.

**Tab. S2 Analysis of lipids identified by MS from eluates of SGs.** Data from Tab. S1 were analyzed for mean (mol%) ± SD for each lipid class (first tab), mean number of double bonds ± SD (second tab) and mean acyl chain length ± SD (third tab).

**Tab. S3 Proteins identified by MS from eluates of SGs.** Cells were labeled with (plus) or without (minus) BC-Fluo and SGs purified. The data from four biological replicates, which were measured in quadruplicates, were log transformed and filtered for contaminants and a coverage of 75%.

**Tab. S4 Significance analysis of identified proteins.** Data from Tab. S3 were analyzed for significant enrichment of proteins from SGs of cells labeled with (plus, first tab) or without (minus, second tab) Fluo. Significance was calculated using permutation-based t-tests with an FDR of 0.05 and S0 of 0.1

**Tab. S5 Proteins identified by MS from eluates of young and old SGs.** Cells were labeled with (plus) or without (minus) BC-Fluo for young and old SGs and purified. The data from four biological replicates were log transformed and filtered for contaminants and a coverage of 75%.

**Tab. S6 Significance analysis of identified proteins.** Data from Tab. S5 were analyzed for significant enrichment of proteins from SGs of cells labeled for young (first tab) or old (second tab) SGs. Significance was calculated using permutation-based t-tests with an FDR of 0.05 and S0 of 0.1

**Tab. S7 Proteins identified by MS exclusive for eluates of young and old SGs.** Data from Tab. S6. Shown are the proteins significantly enriched and exclusively found in eluates of young (first tab) or old (second tab) SGs.

## Material and methods

### Cell and islet culture

The phogrin-CLIP line was cultured in standard INS-1 medium as described before (Ivanova et al., 2013; Hoboth et al., 2015). To maintain the stable integration of phogrin-CLIP, the medium was supplemented with 350 µg/ml G-418.

### Cloning of phogrin-CLIP

Mouse phogrin was obtained from the IMAGE consortium (clone BC_133678) and CLIPf from NEB. The fusion protein was generated by standard molecular cloning techniques. Phogrin was mutated (C931S) as described earlier to abolish phosphatase activity (Caromile et al., 2010). We then connected phogrin and CLIPf with a HRV 3C protease cleavage site (LEVLFQGP) flanked by linkers on each side.

### Generation of BC-Fluorescein(piv)_2_

Solvents for chromatography and reactions were purchased HPLC grade (Sigma-Aldrich, 99.8%, extra dry over molecular sieves). If necessary, solvents were degassed either by freeze-pump-thaw or by bubbling N_2_ through the vigorously stirred solution for several minutes. Unless otherwise stated, all other reagents were used without further purification from commercial sources.

LC-MS was performed on a Shimadzu MS2020 connected to a Nexera UHPLC system equipped with a Waters ACQUITY UPLC BEH C18 (1.7 μm, 50 × 2.1 mm). Buffer A: 0.1% FA in H_2_O Buffer B: acetonitrile. The typical gradient was from 10% B for 0.5 min gradient to 90% B over 4.5 min 90% B for 0.5 min gradient to 99% B over 0.5 min with 1 ml/min flow. Retention times (t_R_) are given in minutes (min).

Preparative and analytical RP-HPLC was performed on a Waters e2695 system equipped with a 2998 PDA detector for product collection (at 230 nm) on a Supelco Ascentis® C18 HPLC Column (preparative: 5 μm, 250 × 21.2 mm; analytical: 5 μm, 250 × 10 mm). Buffer A: 0.1% TFA in H_2_O Buffer B: acetonitrile. The typical gradient was from 10% B for 5 min gradient to 90% B over 45 min 90% B for 5 min gradient to 99% B over 5 min with 8 ml/min flow (preparative) or 4 ml/min (analytical).

High resolution mass spectrometry was performed using a Bruker maXis II ETD hyphenated with a Shimadzu Nexera system. The instruments were controlled *via* Brukers otofControl 4.1 and Hystar 4.1 SR2 (4.1.31.1) software. The acquisition rate was set to 3 Hz and the following source parameters were used for positive mode electrospray ionization: End plate offset = 500 V; capillary voltage = 3800 V; nebulizer gas pressure = 45 psi; dry gas flow = 10 l/min; dry temperature = 250°C. Transfer, quadrupole and collision cell settings are mass range dependent and were fine-adjusted with consideration of the respective analyte’s molecular weight. For internal calibration sodium formate clusters were used. Samples were desalted *via* fast liquid chromatography. A Supelco Titan^TM^ C18 UHPLC Column, 1.9 µm, 80 Å pore size, 20 × 2.1 mm and a 2 min gradient from 10% to 98% aqueous MeCN with 0.1% FA (H_2_O: Carl Roth GmbH + Co. KG ROTISOLV® Ultra LC-MS; MeCN: Merck KGaA LiChrosolv® Acetonitrile hypergrade for LC-MS; FA - Merck KGaA LiChropur® Formic acid 98%-100% for LC-MS) was used for separation. Sample dilution in 10% aqueous MeCN (hypergrade) and injection volumes were chosen dependent of the analyte’s ionization efficiency. Hence, on-column loadings resulted between 0.25–5.0 ng. Automated internal re-calibration and data analysis of the recorded spectra were performed with Bruker’s DataAnalysis 4.4 SR1 software.

In an Eppendorf tube, 1.0 equiv. of isomerically pure 3-oxo-3’,6’-bis(pivaloyloxy)-3H- spiro[isobenzofuran-1,9’-xanthene]-6-carboxylic acid (20.0 mg, 36.7 μmol) was dissolved in DMSO (500 μl) and 4.0 equiv. of DIPEA (19.0 mg, 147 μmol, 24.3 μl) before TSTU (13.3 mg, 44.1 μmol, 1.2 equiv.) was added in one portion. The mixture was allowed to incubate for 10 minutes before 1.5 equiv. of BC-NH_2_ (12.7 mg, 55.1 μmol) were added. The mixture was vortexed and allowed to incubate for 3 h at RT before it was quenched by addition of 20 equiv. of acetic acid and 25 vol% of water. HPLC (MeCN:H_2_O+0.1% TFA = 30:70 to 90:10 over 60 minutes) provided 17.0 mg (22.5 μmol) of the desired compound as white powder in 61% yield.

### Age-dependent labeling of SNAP- and CLIP-tagged proteins

Both Insulin-SNAP and phogrin-CLIP were labeled as previously described (Ivanova et al., 2013; Hoboth et al., 2015). Briefly, INS-1 medium was changed for medium containing either non-fluorescent 5 µM CLIP/SNAP-Cell Block, 2 µM SNAP-Cell^®^ 430, 0.6 µM CLIP-Cell TMR-Star^®^ or 0.6 µM synthesized CLIP-Fluorescein as indicated in the text. After 30 min the medium was aspirated, the cells washed twice with PBS and replaced by non-modified medium. The cells were washed twice for 30 min and once for 60 min.

### Sample preparation for fluorescence microscopy, image acquisition and analysis

For SIM imaging, phogrin-CLIP cells were grown on high precision coverslips coated with poly-ornithine until a confluency of about 70-80%. The cells were then labeled with the respective SNAP/CLIP substrates as indicated above. The cells were fixed with 4% PFA for 20 min and permeabilized with 0.1% Triton X-100 for 10 min. The permeabilized cells were blocked for 1 hour with 0.2% (fish skin) gelatine solution + 0.5% BSA in PBS at RT. The primary antibodies were diluted in blocking solution and the cells labeled for 30 min at RT. Secondary antibody staining was performed similarly for 20 min. For staining of EEA1 (Thermo Scientific, MA5-14794) and LAMP2 (Thermo Scientific, PA1-655) permeabilization was ensured by the presence of 0.1% saponin. Anti-insulin was from Sigma (I-2018) and anti-SNAP from NEB (P9310S). The coverslips were mounted on glass slides with VectaShield Antifade Mounting Medium and fixed with nail polish. Images were acquired using a DeltaVision OMX SIM with an Olympus Plan ApochromatN 60x oil objective with a NA of 1.42. Stacks with a z step-size of 125 nm were acquired and reconstructed with the SoftWoRx software package (SoftWoRx, Germiston, South Africa). Images were further processed and analyzed using FIJI and R (R Core Team, 2020; Wickham, 2016). Contrast was changed for representation purposes. Quantification was performed on background-subtracted raw images.

### Insulin SG purification for SDS-PAGE and Western Blot

For isolation of insulin SGs, phogrin-CLIP cells were grown in 175 cm^2^ flasks (typically 3-4 per condition). After labeling as indicated above, the cells were washed once with PBS, then harvested in 10 ml PBS per flask, spun down for 5 min at 1200 rpm, 4°C and the cell pellets resuspended in 500 µl homogenization buffer (250 mM sucrose, 150 mM NaCl, 4 mM HEPES pH 7.4, 1 mM EGTA, 1% Protease Inhibitor Cocktail). For lipidomic and proteomic analyses, 1.5 mM Na_3_VO_4_ and 1 µM PI-PLC inhibitor U73122 (Tocris) were added to the homogenization buffer. The cell suspensions were homogenized with 12 strokes of a glass homogenizer on ice and the homogenates spun down for 2 min at 1200 rpm, 4°C to remove nuclei and cell debris. From this point on, flasks may be pooled and numbers indicate amounts/volumes per 175 cm^2^ flask. Aliquots of the postnuclear supernatants (input, 20 µl) were lysed with lysis buffer and measured by BCA to adjust protein contents of different treatments. Of each condition 10 µg protein were aliquoted for quality control. Anti-substrate antibodies (5 µg per flask) were added to the cell homogenate and rotated for 2 h at 4°C (9 rpm). For each flask, 30 µl Protein G Dynabeads (ThermoFisher) were washed once in 150 µl PBS-T 0.02% and washed another two times with 150 µl HB. The beads were resuspended in 50 µl homogenization buffer and added to the homogenate-antibody suspension and rotated for 1 hour at 4°C (9 rpm). The suspension-containing tubes were placed in a magnetic rack and the supernatant transferred in a new tube (flow-through). The same volume as for the input was taken from the flow-through for quality control. The beads were washed three times in 300 µl homogenization buffer per flask for 30 min at 4°C (9 rpm). For three 175 cm2 flasks, the beads were resuspended in 400 µl homogenization buffer and 10 µg HRV 3C protease added and rotated over night. The suspension was placed on a magnetic rack and let sit for 30-60 s and the supernatants were taken and placed in a new tube in the magnetic rack to remove residual magnetic beads. This step was repeated once. The beads were boiled in SDS-loading buffer for analysis by Western Blot. To remove potential lysosomal organelles, 2 µg/flask of LAMP2 antibody (Thermo Scientific, 51-2200 or self-made) were added to the eluates and rotated for 3 h at 4°C (9 rpm). For each three flasks 50 µl of Protein G Dynabeads were prepared as above and the eluates transferred to the bead-containing tubes. The tubes were then rotated for 1 h at 4°C (9 rpm), placed on magnetic racks and the eluates transferred to a new tube in the magnetic rack. This step was repeated once and the resulting eluate could be used for Western Blot or other analyses.

Antibodies against ICA512 were self-raised, those against phogrin were a kind gift from Howard Davidson. Anti-SNAP was from NEB (P9310S), anti-CHGA from Abcam (ab45179), anti-PDI from Stressgen (SPA-891), anti-GST from Santa Cruz (sc-138), anti-HIS from Novagen (70796-3), anti-TGN38 from BD Transduction (610898), anti-PC2 from GeneTex (GTX114625), anti-CPE from Sigma (AB5314), anti-γ-tubulin from Sigma (T- 6557), anti-CANX from BD Transduction (610523), anti-ICA69 from Abcam (ab81500), anti- GM130 from BD Transduction (610822) and anti-SYP1 from Synaptic Systems (101011), anti-EEA1 (Thermo Scientific, MA5-14794), anti-LAMP2 (Thermo Scientific, 51-2200).

### Real-time PCR

Real-time PCR was performed using the GoTaq qPCR Master Mix (Promega) according to manufacturer’s instructions. Briefly, cDNA from RT reactions was diluted 1:2 in RNase-free water. Triplicate reactions were set up in Semi-skirted 96-Well PCR Plates (0.2 ml) with optical strip caps (Agilent). The PCR reactions were carried out in an AriaMx Real-time PCR System (Agilent). For absolute quantification, serial dilutions of the target sequence cloned into pCRII vectors were used. The results were then normalized by parallel amplification of rat β-actin mRNA. The following primers were used for the detection of rat β-actin (fwd: 5’-CAA CGG CTC CGG CAT GTG CAA GG-3’; rev: 5’-TCT TCT CCA TAT CGT CCC AGT TG-3’), γ-tubulin (fwd: 5’-CAA CAG TCC TGG ATG TCA TGA GG-3’; rev: 5’-GGT GTG GTT GGC CAT CAT GAG C-3’), phogrin (fwd: 5’-TCC AGA CAA AGG AGC AGT TT-3’; rev: 5’-GAG TCT GAA GGA CCC CCT TA-3’) and phogrin-CLIP (fwd: 5’- CAC TCC CAC TGG AGG TTT TA-3’; rev: 5’-ACC CAG ACA GTT CCA GCT T-3’).

### Insulin secretion

Static insulin secretion was measured using an ultrasensitive insulin HTRF kit (Cisbio) according to manufacturer’s recommendations.

### Sample preparation for electron microscopy

For electron microscopy, insulin SGs were purified as described above. Instead of acetone precipitation, SGs were either used directly for negative staining or fixed for epon embedding.

For negative staining of purified granules 5-10 ml of sample was pipetted to a 300-mesh copper grid covered with a thin carbon-coated and glow discharged formvar film, and incubated for 10 min to allow granules to sediment and adhere to the film. Liquid was removed with a piece of filter paper, shortly washed with a drop of water (2x) and stained with 1% uranyl acetate (UA) in water for about 20 sec. UA was removed slowly with filter paper and the grid was air dried before inspection.

Protein G bead bound granules were fixed with a mixture of formaldehyde (FA, prepared from paraformaldehyde prills) and glutaraldehyde (GA) (2% FA/2% GA in 100 mM phosphate buffer), centrifuged, and resuspended in lukewarm agarose (2%). The agarose was cooled down and cut into small blocks for further processing. Samples were postfixed in 2% aqueous OsO_4_ solution containing 1.5% potassium ferrocyanide and 2 mM CaCl_2_. After washes in water, samples were incubated in 1% thiocarbohydrazide, washed again and contrasted in 2% osmium in water for a second time (Hanker et al., 1966). Samples were washed in water, *en bloc* contrasted with 1% uranyl acetate/water, washed again in water, dehydrated in a graded series of ethanol, infiltrated in the epon substitute EMbed 812 (1+2, 1+1, 2+1 epon/ethanol mixtures, 2x pure epon), and finally embedded in flat embedding molds. Samples were cured at 65°C in the oven overnight, and ultrathin sections were cut with a Leica UC6 ultramicrotome and collected on formvar-coated slot grids. Sections were contrasted with uranyl acetate and with lead citrate (Venable and Coggeshall, 1965). All samples were imaged either with a FEI Morgagni 268D (Thermo Fisher Scientific, equipped with a Megaview III camera, SIS-Olympus) or with a Jeol JEM1400 Plus (equipped with a Ruby camera, JEOL) both running at 80 kV acceleration voltage.

### Sample preparation for shotgun lipidomics

Extraction of lipids was performed with a modified version of a previously reported procedure (Ejsing et al., 2009, Sampaio et al., 2011). Granules were extracted for 15 min on ice, using 10 volumes of CHCl_3_/MeOH (10:1 v/v) including the IS mixture used for absolute quantification. Samples were centrifuged (6000xg, 5min, 4°C), the organic phase was collected and the water phase was re-extracted with 8 volumes of CHCl_3_/MeOH/acetone/1M HCl (2:1:0.5:0.1 v/v) for 10 min on ice with thorough vortexing. Samples were centrifuged (6000xg, 5min, 4°C) and the organic phase was collected, pooled and afterwards evaporated under a nitrogen stream for additional 6 hours. The sample was re-suspended in 30 – 50 µl CHCl3/MeOH (1:2 v/v) for MS analysis. The samples were kept at 4°C during the whole sample preparation and extraction procedure in order to prevent lipid degradation.

### Annotation of lipid species and standard mixture

Glycerophospholipids, DAG and TAG species were annotated as: (lipid class) (number of carbons in all fatty acids)-(number of double bonds in all fatty acids).

The internal (IS) standard mixture contained the following: 40 pmol of 1-heptadecanoyl-2- (5Z,8Z,11Z,14Z-eicosatetraenoyl)-sn-glycero-3-phosphocholine (PC 17:0-20:4), 20 pmol of 1-tridecanoyl-sn-glycero-3-phosphocholine (LPC 13:0), 35 pmol of 1-heptadecanoyl-2- (5Z,8Z,11Z,14Z-eicosatetraenoyl)-sn-glycero-3-phospho-L-serine (PS 17:0-20:4), 10 pmol of 1-heptadecanoyl-2-(5Z,8Z,11Z,14Z-eicosatetraenoyl)-sn-glycero-3-phospho-(1’-rac-glycerol) (PG 17:0-20:4), 20 pmol of 1-heptadecanoyl-2-(5Z,8Z,11Z,14Z-eicosatetraenoyl)- sn-glycero-3-phosphate (PA 17:0-20:4), 40 pmol of d5-Diglyceride (DAG D5), 30 pmol of N-(dodecanoyl)-sphing-4-enine-1-phosphocholine (SM C12), 40 pmol of 1-heptadecanoyl- 2-(5Z,8Z,11Z,14Z-eicosatetraenoyl)-sn-glycero-3-phosphoethanolamine (PE 17:0-20:4), 30 pmol of N-(dodecanoyl)-sphing-4-enine (CER C12).

### Mass spectrometry of lipids

Lipid extract was mixed 1:1 with 7.5 mM ammonium formate solution (dissolved in CHCl_3_/MeOH/*i*-ProOH 14:28:58 v/v). For each analysis, 30 µl of samples were loaded onto 96-well plates (Eppendorf, Hamburg) and sealed with aluminum foil. Mass spectrometry analysis was performed on a QExactive instrument equipped with a robotic nanoflow ion source TriVersa NanoMate (Advion Biosciences) using nanoelectrospray chips with a spraying nozzle diameter of 4.1 µm. The ion source was controlled by the Chipsoft 8.3.1 software. The temperature of the ion transfer capillary was 25°C; S-lens RF level was set to 50%. Extract 1 and 2 (10 µl each) were analyzed either in positive and negative ion mode for 1.5 min in a single acquisition at a resolution of R_m/z=200_=140,000 for FT-MS or in positive and negative ion mode for 38 min in a single acquisition at a resolution of R_m/z=200_=140,000 for FT-MS and R_m/z=200_=70,000 for FT-MS/MS. Samples were infused with a backpressure of 1.25 psi and spray voltage of +0.96 kV and -0.96 kV, respectively. In order to avoid initial spray instability, the delivery time was set to 30 sec. Polarity switch from positive to negative ion mode was set at 0.5 resp. 16.5 min after contact closure, followed by a lag of 20 sec resp. 30 sec after polarity switch for spray stabilization. FT-MS acquisition method starts with positive ion mode for 0.5 min by acquiring the *m/z* 400-1600 using automated gain control (AGC) of 3 x 10^6^ and maximum ion injection time (IT) of 500 ms. FT-MS/MS+ experiments were triggered by an inclusion list for 16 min, including all masses from 400.20 to 1,000.86 with 1 Da intervals and a normalized collision energy (NCE) of 15%. AGC and maximum IT were set to 2 x 10^4^ and 650 ms, respectively. FTMS acquisition in negative ion mode was set for 0.5 min by acquiring the *m/z* 400-1,600 using AGC of 3 x 10^6^ and IT of 500 ms. FT-MS/MS- experiments were triggered again by an inclusion list for 20.5 min, including all masses from 400.20 to 1,000.86 with 1 Da intervals and NCE of 23%, AGC of 2 x 10^4^ and maximum IT of 650 ms. Target masses for MS/MS were specified in the inclusion list. Raw data are available in Tab. S1.

### Lipid identification and quantification/Data processing and Analysis

Data were analyzed with an in-house developed lipid identification software based on LipidXplorer (Herzog et al., 2011; Herzog et al., 2012). Molecular Fragmentation Query Language (MFQL) queries were compiled for all lipid classes included in the IS mix. Identification was based on combining MS precursor (mass accuracy better than 5 ppm) and MS/MS fragmentation. Data post-processing and normalization were performed using an in-house developed data management system. Only lipid identifications with a signal-to-noise ratio >5 and a signal intensity 5-fold higher than in corresponding blank samples were considered for further data analysis. All downstream analyses were performed in R (R Core Team, 2020). Only the lipids measured in 75% or more of the samples were kept. The dataset was log2-scaled before Principal Component Analysis. The descriptive analysis was performed on the mol %-transformed dataset, i.e. picomole quantities were divided by the sum of the lipids detected in the respective sample and multiplied by 100. Total carbon chain length and double bonds plots result from grouping together all the lipids that present the same number of carbon atoms (total length) or the same number of double bonds (degree of unsaturation) and calculating the mean and standard deviation in each group of samples (young, old and mixed). Plots were generated using ggplot2 (Wickham, 2016).

### Lipid extraction and GP measurements

For characterization of the biophysical properties of the lipids, granule samples were extracted using a two-step extraction protocol (chloroform:methanol 10:1 followed by 2:1 (Ejsing et al., 2009). After each step, the lipid containing organic phase was pooled and dried under a nitrogen stream followed by incubation undera vacuum for 4 h to remove organic solvents. The dried lipid film was re-hydrated in 100μl of water, for 15 in at 600 rpm. To yield unilamellar vesicles (LUVs), samples were subjected to 10 cycles of freeezing in liquid nitrogen and subsequent thawing in a heating block at 40°C. The vesicle solution was extruded 21 times through a 100 nm diameter polycarbonate membrane (Whatman® Nuclepore, Fisher Scientific, US) using an extrusion kit (Avanti, US). Membrane order of liposomes prepared form granule lipids was assessed by determining C-laurdan generalized polarization (GP) indices. Liposomes were stained with C-laurdan (Kim et al., 2007) and incubated for 30 min in the dark at RT. Fluorescence spectra were obtained with a FluoroMax-4 spectrofluorometer (Horiba) equipped with a temperature-controlled Peltier element (Newport) at 23°C. Excitation was 385 nm and emission recorded at 400-600 nm with 1 nm. Spectra were background-corrected and GP values were calculated as described in Kaiser et al., (2009).

### Sample preparation - proteomics

Samples used for proteomics analysis were reconstituted in 500 µl 6M GdmCl, 100 mM Tris-HCl pH 8.5 solubilization buffer. After addition of 10 mM TCEP and 55 mM CAA the samples were boiled for 20 min at 95°C, 800 rpm on a thermoshaker (Eppendorf) followed by sonication at maximum power (Bioruptor) for 10 cycles of 30 seconds of sonication and 30 seconds of cooldown each. The sample was briefly spun down and boiled again for 10 min at 95°C, 800 rpm. 1 ml of 100 mM TrisHCl pH 8.5 was added to each sample to dilute the GdmCl molarity to below 2 M in total. 10 µg of Trypsin and LysC each were added to the sample solution followed by overnight digestion at 37°C, 800 rpm on a thermoshaker (Eppendorf). Next day, the sample was acidified to 1% TFA followed by stage-tip cleanup via styrene-divinylbenzene reverse-phase sulfonate (SDB-RPS). Sample liquid was loaded on two 14-gauge stage-tip plugs fixed within a 200 µl pipette tip. Peptides were cleaned up with 2x 100 µl 99% ddH2O 1% TFA into 2x 100 µl 99% Isopropanol 1% TFA at 1200 xg at RT each using a table-top centrifuge (Eppendorf). Afterwards, peptides were eluted with 80 % acetonitrile, 5% ammonia, 15% ddH_2_O and dried at 30°C in a SpeedVac centrifuge (Eppendorf). Finally, peptides were reconstituted in 4.2 µl of 2% acetonitrile, 0.1% TFA,

### Liquid chromatography and mass spectrometry of proteins

LC–MS was performed with an EASY nanoLC 1200 (Thermo Fisher Scientific) coupled online to a modified trapped ion mobility spectrometry quadrupole time-of-flight mass spectrometer (timsTOF Pro, Bruker Daltonik) via nano-electrospray ion source (Captive spray, Bruker Daltonik; Brunner et al., 2021). Peptides were loaded on a 50 cm in-house-packed HPLC-column (75 µm inner diameter packed with 1.9 µm ReproSil-Pur C18-AQ silica beads, Dr. Maisch). Sample analytes were separated using a linear 120 min gradient from 3% to 30% buffer B in 95 min followed by an increase to 60% for 5 min, and by a 5 min wash at 95% buffer B at 300 nl/min (buffer A: 0.1% formic acid, 99.9% ddH_2_O; buffer B: 0.1% formic acid, 80% CAN, 19.9% ddH_2_O). The column temperature was kept at 60°C by an in-house-manufactured oven.

Data were acquired in data-dependent PASEF mode with 1 MS1 survey TIMS–MS and 10 PASEF MS/MS scans per acquisition cycle. Ion accumulation and ramp time in the dual TIMS analyser was set to 100 ms each and we analyzed the ion mobility range from 1/*K*0 = 1.6 Vs cmm-2 to 0.6 Vs cm-2. Precursor ions for MS/MS analysis were isolated with 2Th windows for *m/z* < 700 and 3Th for *m/z* > 700 in a total *m/z* range of 100–1,700 by synchronizing quadrupole switching events with the precursor elution profile from the TIMS device. The collision energy was lowered linearly as a function of increasing mobility starting from 59 eV at 1/*K*0 = 1.6 VS cm-2 to 20 eV at 1/*K*0 = 0.6 Vs cm-2. Singly charged precursor ions were excluded with a polygon filter (otof control, Bruker Daltonik). Precursors for MS/MS were picked at an intensity threshold of 2,500 a.u. and resequenced until reaching a ‘target value’ of 20,000 a.u., taking into account a dynamic exclusion of 40 seconds of elution. Raw data are deposited on the PRIDE database with identifier XYZ and are available upon request.

### Proteomics raw file processing

Raw files were searched against the rat UniProt databases UP000002494_10116.fa and UP000002494_10116_additional.fa with MaxQuant (v.1.6.7), which extracts features from four-dimensional isotope patterns and associated MS/MS spectra (Prianichnikov et al., 2020). FDRs were controlled at 1% both on peptide spectral match (PSM) and protein level. Peptides with a minimum length of seven amino acids were considered for the search including *N*-terminal acetylation and methionine oxidation as variable modifications and cysteine carbamidomethylation as fixed modification, while limiting the maximum peptide mass to 4,600 Da. Enzyme specificity was set to trypsin cleaving carboxy terminal to arginine and lysine. A maximum of two missed cleavages were allowed. Maximum precursor and fragment ion mass tolerance were searched as default for TIMS-DDA data, while the main search peptide tolerance was set to 20 ppm. The median absolute mass deviation for the data was less than one ppm. Label-free quantification was performed with the MaxLFQ algorithm and a minimum ratio count of one (Cox et al., 2014).

### Bioinformatic analysis of proteomics data

Proteomic data was analyzed using Perseus (v.1.6.7.0). Reverse database, contaminant, and only by site modification identifications were removed from the data set. Median values were calculated for technical replicates and log2-transformed. The data were then filtered for at least 75% completeness per group. Missing values were then imputed from a data table specific normal distribution estimate with a downshift of 1.8 and a width of 0.3 SD. Samples of unlabeled and labeled SGs, as well as young and old SGs were tested for differences in their medians using a permutation-based two-sided Student’s t-test with 250 iterations, a FDR of 0.05 and a S0 of 0.1. The data was presented as volcano plots. Principal component analysis was performed for samples of young and old SGs to visualize data reproducibility and variability.

